# Material-specific quarantine durations for SARS-CoV-2 inactivation on musical instruments and music-related materials

**DOI:** 10.64898/2026.07.01.735763

**Authors:** Pastorino Boris, Touret Franck, Viala Romain, Creton Milena, Morand Jean Charles, Billecard François, Reyre Fanny, Jousserand Michael, N. Charrel Remi

**Author notes:** Corresponding author: email: Boris Pastorino. Franck Touret, Romain Viala, Milena Creton, Jean Charles Morand, Fanny Reyre, Michael Jousserand, François Billecard, Remi N. Charrel.

## Abstract

The COVID-19 pandemic has imposed a reevaluation of safety protocols across various sectors, including the arts. This study addresses a critical gap in understanding SARS-CoV-2 persistence on materials commonly associated with musical instruments and scores, such as alloys, varnishes, reeds, and paper. While previous research has explored viral survival on various surfaces, limited data exists for materials specific to musical contexts. In this work, we investigate the efficacy of quarantine as a non-destructive method for inactivating SARS-CoV-2 on 16 materials, including brass, silver plating, ABS plastic, ebonite, and various varnishes and paper types.

Results revealed significant variability in viral persistence across materials. Non-porous surfaces like metals and ABS plastic cleared infectivity within 3 days, while porous materials such as reeds and music scores required up to 7 days. Gold-plated brass and certain varnishes showed intermediate persistence, with infectivity clearing after 4 days. These findings are in agreement with prior studies indicating that SARS-CoV-2 survival is highly dependent on surface composition, with porous and organic-coated materials retaining viable virus longer due to reduced environmental stress.

Our results highlight the feasibility of stratified quarantine protocols based on material type, offering practical guidelines for musicians and institutions and provides critical insights for mitigating SARS-CoV-2 transmission risks in musical settings.

## INTRODUCTION

The emergence of SARS-CoV-2, an airborne transmitted coronavirus [1], and the subsequent global health crisis have imposed a reevaluation of safety protocols across various domains, including the arts and cultural sectors. Notable outbreaks of SARS-Cov-2 have occurred at mass gatherings associated with artistic activities, particularly music concerts [2]. Among these, the handling of musical instruments and related materials has posed unique challenges for both professionals and amateurs to prevent SARS-CoV-2 transmission. The porous and complex nature of many musical instruments, combined with frequent physical contact, including blowing, raised concerns about potential viral transmission [3, 4]. In addition, it was shown that playing a wined instrument produces more aerosol emissions than talking or breathing in a quiet way [4]. As a result, there was an urgent need to establish safe practices that allow musicians to continue their professional or recreative activities while minimizing the risk of infection for both performers and audiences [5-7].

This study addressed a critical gap in the understanding of SARS-CoV-2 persistence on materials commonly associated with musical instruments and scores. While previous research has explored viral survival on various surfaces, there was limited data specifically tailored to the diverse materials found in musical contexts, such as alloys, varnishes, reeds, and paper [8, 9]. The goal of this study was to investigate the efficacy of quarantine as a non-destructive method for inactivating SARS-CoV-2 on these materials. By determining the duration required for viral inactivation under controlled conditions, we aimed to provide practical guidelines that can be implemented in real-world settings, such as orchestras, music schools, and individual practice environments.

We focused on 16 materials commonly found in musical instruments and scores, including various alloys, varnishes, and types of paper. Experiments were conducted under controlled environmental conditions (19-21°C and 50-60% relative humidity) to simulate typical indoor settings. The study consisted of two phases: the first phase evaluated the effect of a 72-hour quarantine on viral infectivity, while the second phase explored shorter or longer quarantine durations based on the initial results. By adopting a material-specific approach, this research has attempted to provide differentiated recommendations that take into account the unique characteristics of each surface.

## MATERIALS AND METHODS

### Virus Strain and Titration

Experiments were performed in BSL3 facilities located in the UVE premises at the School of Medicine using a clinical SARS-CoV-2 strain UVE/SARS-CoV-2/2020/FR/702) (https://www.european-virus-archive.com/virus/sars-cov-2-virus-strain-uvesars-cov-22020fr702). The strain was inoculated at a 0.001 MOI in 90% confluent Vero-E6 cells (ATCC number CRL-1586) and incubated at 37°C for 48 h. The supernatant was collected, clarified by spinning at 1500 g for 10 min, supplemented with 25mM HEPES (Sigma-Aldrich, Lyon, France), and aliquots were stored at −80°C. One aliquot was thawed and used for titration using 50% tissue culture infectivity dose (TCID50); briefly, when cells were at 90% confluence, six replicates were infected with 150µL of ten-fold serial dilutions of the virus aliquot and incubated for 4 days at 37°C under 5% CO2. Cytopathic effect (CPE) was read using an inverted microscope, and infectivity was expressed as TCID50/mL using the Karber formula [10]. All samples were quantified by end-point titration on Vero E6 cells with a limit of detection (LoD) of ^100.5^ TCID50/mL (0.8 log10TCID50/mL).

### Studied materials

Based on the advices provided by some of us (RV, MC, JCM, FR, MJ and FB) who have considerable expertise and experience in music instruments field, five alloys were selected: Brass 70/30, German Silver (also known as Silver Nickel), Gold plating, Nickel plating, Silver plating. Other materials components of music instruments such as ebonite, ABS or reed were selected. Five types of varnish (polyurethane, nitrocellulose, shellack, linseed oil, epoxyde) were included. Finally, three different types of paper used for music scores were selected.

### Protocol Phase 1

Briefly, a 50 μL droplet of virus culture (∼5·8 log unit of TCID_50_ per mL) was pipetted on various surfaces in triplicate (∼1cm^2^ per piece) and the inoculated material was stored at 19-21°C (50-60% relative humidity) in a BSL-3 laboratory for quarantine times varying from 1 day to 7 days. After contact, SARS-CoV-2 was recovered by pipetting on the surface for a minute, and collecting as much as possible of the fluid into a microtube before virus titration (LoD is 0.8 log10TCID50/mL). All conditions were tested in triplicate and results are presented as mean value.

Whether there was no residual infectivity after 3-day quarantine, then quarantine times of 1 and 2 days were tested in a second protocol phase (see below). Whether residual infectivity was observed after a 3-day quarantine, then quarantine times ranging from 4 to 7-day were tested (Fig.1).

**Figure 1.**
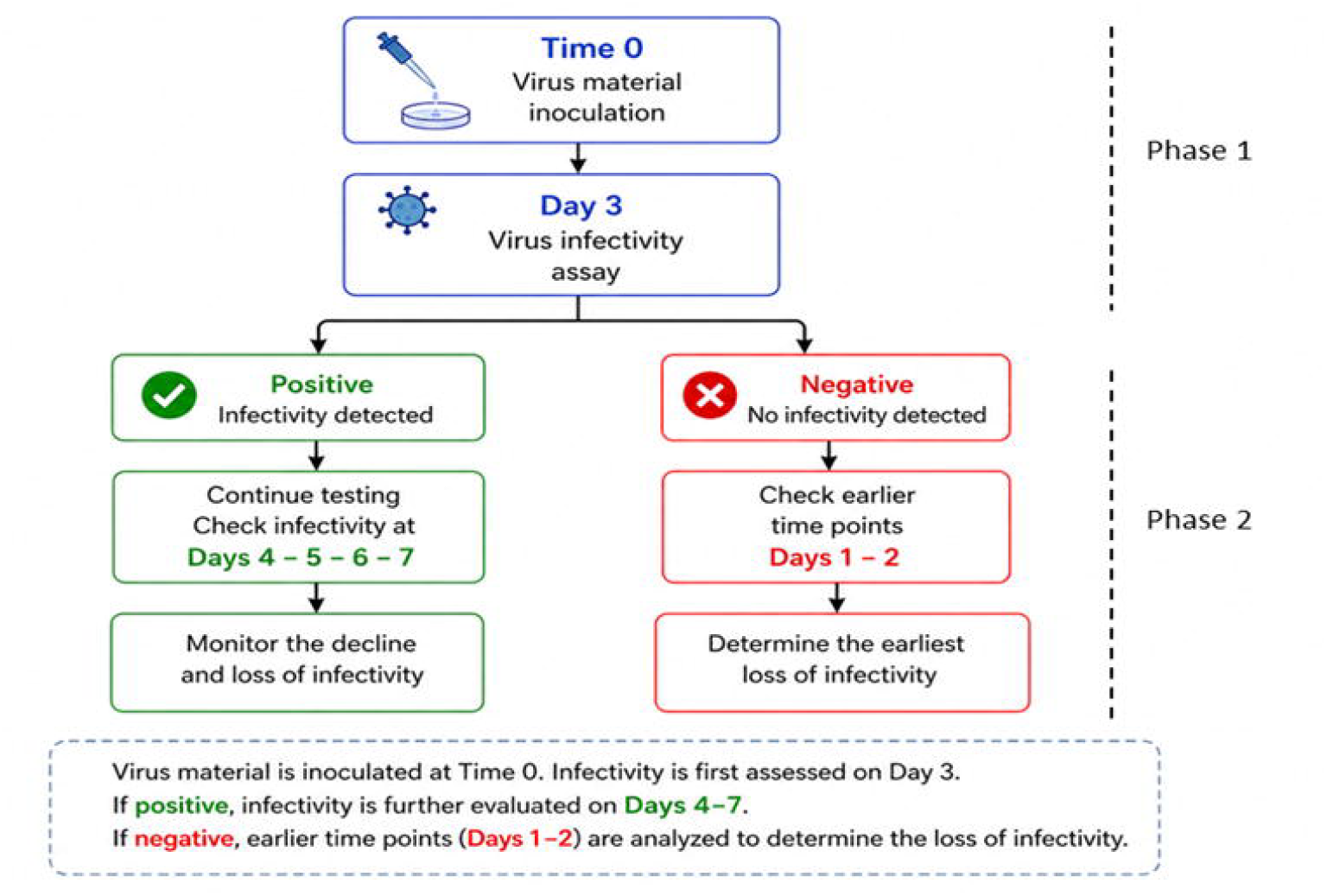
Workflow for evaluating the persistence of SARS-CoV-2 infectivity on instruments and related materials.

### Protocol Phase 2

The protocol was similar to that described in Phase 1. The technical aspects and the inoculum were identical. Instead of titration after SARS-CoV-2 recovery, cells were inoculated and incubated at 37 °C under 5% CO2 for 5 days. The read-out was the presence or absence of cytopathic effect (CPE) after observation with an inverted microscope. Compared with Phase 1, Phase 2 read-out was not quantitative but qualitative. All conditions were tested in triplicate. Results were analyzed as presence (+) or absence (-) of an observed CPE (Fig.1).

### Statistical Analysis

The initial inoculum was used as a reference to calculate log_10_ reduction factors. Values below the assay LoD (0.8 log_10_ TCID_50_/mL) were set conservatively at 0.8 log_10_ TCID_50_/mL for statistical analysis. The mean, standard deviation (SD), and approximate 95% confidence interval (CI) of log_10_ TCID_50_/ml were calculated for each material from three independent replicates, assuming a normal distribution. Due to the small sample size (n = 3), the 95% confidence intervals were estimated using the t-distribution with two degrees of freedom.

Because several measurements were at the LoD and the number of replicates per condition was low, no formal hypothesis testing was performed. Instead, differences between materials were interpreted descriptively, considering reduction of at least one log_10_ as potentially meaningful in term of viral infectivity.

## RESULTS

### The two-phase strategy is presented in the figure 1

#### Phase 1: effect of 3-day quarantine

To evaluate whether there are substantial differences in SARS-CoV-2 infectivity and survival, we deposited on various surfaces related to musical instruments or practice, a fixed dose of infectious viral supernatant. We used a protocol already validated for SARS-CoV-2 infectivity evaluation [11]. After 72-hr of quarantine we observed differences in the infectivity of SARS-CoV-2 depending on the type of surface and subsequently on the corresponding instrument (Table 1).

**Table 1.** Effect of 3-day quarantine on SARS-CoV-2 infectivity.

| Category | Material | Mean titer (log<sub>10</sub> TCID<sub>50</sub>/ml) | SD | 95% CI | Mean log reduction |
| --- | --- | --- | --- | --- | --- |
| Metal | Silver plating | 0.8 | 0.0 | 0.8–0.8 | 5.0 |
|  | Nickel plating | 0.8 | 0.0 | 0.8–0.8 | 5.0 |
|  | Gold-plating | 1.5 | 1.16 | 0.3–2.7 | 4.3 |
|  | Nickel silver | 0.8 | 0.0 | 0.8–0.8 | 5.0 |
|  | Brass | 0.8 | 0.0 | 0.8–0.8 | 5.0 |
| Plastic | ABS Plastic | 0.8 | 0.0 | 0.8–0.8 | 5.0 |
|  | Ebonite | 2.8 | 0.0 | 2.8–2.8 | 3.0 |
| Varnish | Polyurethane | 0.8 | 0.0 | 0.8–0.8 | 5.0 |
|  | Nitrocellulose | 0.8 | 0.0 | 0.8–0.8 | 5.0 |
|  | linseed oil | 1.7 | 1.02 | 0.6–2.8 | 4.1 |
|  | Shellac | 0.9 | 0.17 | 0.6–1.1 | 4.9 |
|  | Epoxyde | 1.6 | 1.00 | 0.5–2.7 | 4.2 |
| Organic | Reed | 3.9 | 0.0 | 3.9–3.9 | 1.9 |
|  | Music score 80g copy paper | 4.0 | 0.0 | 4.0–4.0 | 1.8 |
|  | Music score Barenreiter paper | 4.3 | 0.0 | 4.3–4.3 | 1.5 |
|  | Music score ancient paper | Technically impossible* |  |  |  |

\*, technically unsuitable due to excessive porosity. initial inoculum: 5.8 log<sub>10</sub> TCID<sub>50</sub>/ml. Mean titer was calculated from three replicates (n=3). Values below the LoD (<0.8) were conservatively set to 0.8 for analysis. Shaded lines correspond to material for which residual infectivity of SARS-CoV-2 was observed.

Of the five metallic materials, four had cleared infectivity. with 95% confidence intervals (CIs) grouped near the LoD (≤0.8 log_10_ TCID_50_/mL), indicating a reproducible reduction of more than five logs compared to the initial inoculum; only Gold plating showed virus recovery at a titer of 1.5 log_10_ TCID_50_/mL (0.3–2.7; 95% CI) despite a 4.3 log reduction compared with the inoculum.

Regarding plastic-based materials, ABS plastic cleared infectivity whereas ebonite preserved infectious viral particles after a 3-day quarantine.

Of the five varnished materials, polyurethane and nitrocellulose were the only ones to have cleared infectivity. Shellac varnish was very close to complete clearance of infectivity with a titer of 0.9 just above the assay LoD (0.6-1.1; 95% CI). Linseed oil and epoxy varnishes show wider CI values, suggesting intermediate efficacy and less consistency across replicates.

Of the four organic materials, the 3-day quarantine was poorly efficient. CI values centered between 3.9 and 4.3 log_10_ TCID_50_/mL consistent with a much more modest reduction of approximately 1.5 to 1.9 logs. Music scores made on ancient paper were not testable because virus recovery was impossible due to an extreme porosity; therefore they were excluded from the study.

### Phase 2. SARS-CoV-2 infectivity before and beyond 3 days quarantine

A second experiment was carried out, but an extended duration of 7 days, in order to better characterize the exact survival time in relation to a specific material (Figure 2). However, instead of a time-defined titration, we re-cultured the virus every day from Day 3 through Day 7. We used a simpler readout with viral isolation and cytopathic effect observation converted to positive and negative representative residual infectivity (Figure 1). For materials still contaminated after 3 days, residual infectivity was cleared after 4 days for gold-plated brass, ebonite, three varnishes (shellac, linseed oil and epoxyde). Interestingly organic based materials are showing the longest infectivity after 6 days for reed, and after 7 days for music scores.

**Figure 2.**
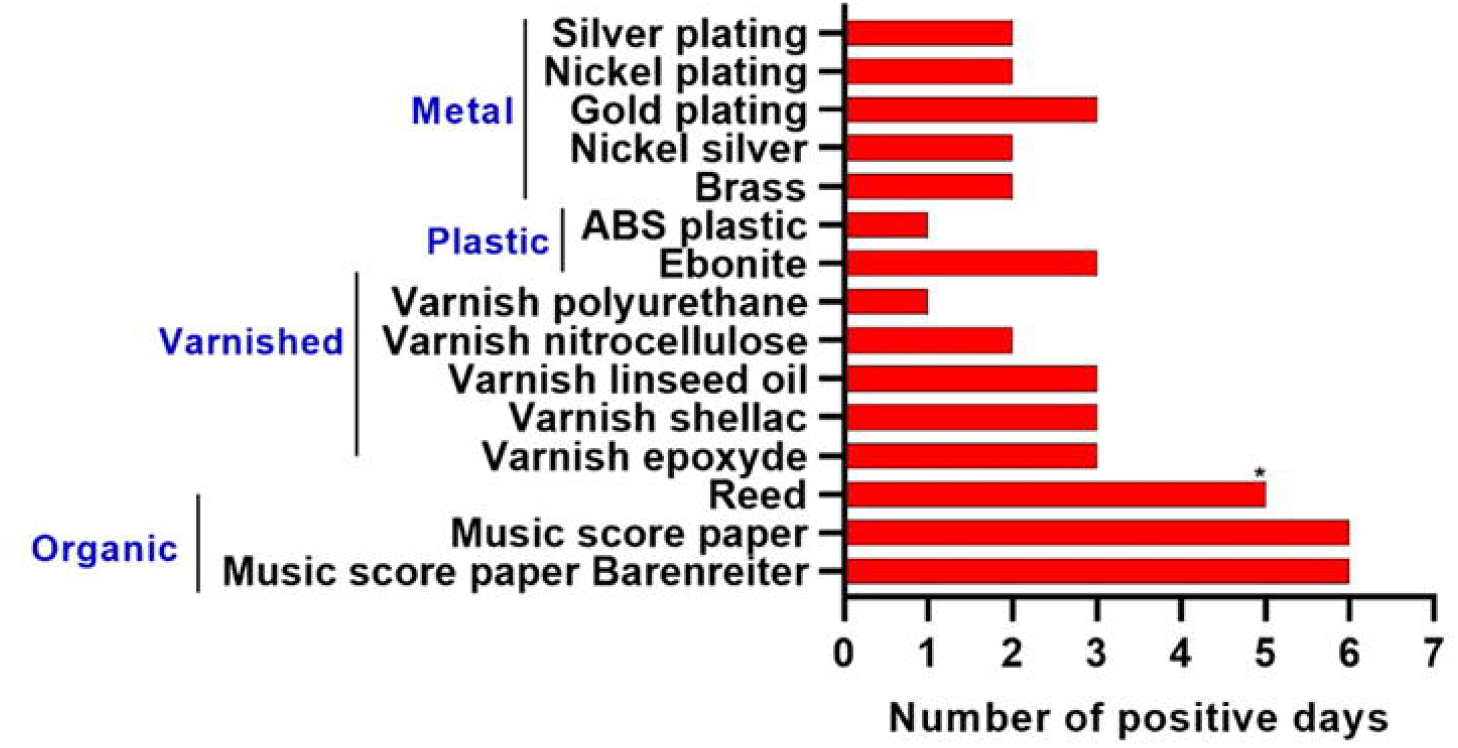
Quarantine duration needed for reducing the SARS-CoV-2 infectiosity below the detection threshold. The red bar represents the number of days for which an infectious title can be measured > 0.8 TCID50/ml. *, not tested since quarantine unsuitable and alternative technique using soaking in RiegerClean solution. Values correspond to residual infectivity as tested in the conditions of the study. All values reflect the mean of 3 replicates.

## DISCUSSION

There is an obvious potential risk that wind and brass instruments, along with singing, can generate aerosols capable of transmitting respiratory viruses such as COVID-19 [5]. Viruses can persist on instrument surfaces, creating a risk of SARS-CoV-2 transmission when instruments or scores are shared, directly or via hand-mediated transmission [3, 12]. Clearly, the combination of a pandemic together with the desire to maintain artistic activities are incentive for developing hygiene protocols in order to suppress or reduce the risk of infection during such activities.

Our study provides critical insights into the feasibility of quarantine as a practical, non-destructive method for inactivating SARS-CoV-2 on materials commonly associated with musical instruments and scores.

Ben-Shmuel et al. (2020) highlighted prolonged viral survival on fibrous and porous materials in quarantine facilities [13]. Similarly, Edwards et al. (2022) underscored material-specific variability in viral persistence on sports equipment [14], suggesting analogous complexities for musical materials. Yet, no study to date has comprehensively evaluated quarantine durations for the specialized alloys, varnishes, and papers ubiquitous in musical practice.

The results demonstrate significant variability in viral persistence across materials, reinforcing the importance of surface-specific inactivation protocols for real-life application. The observed differences in quarantine efficacy align with prior studies showing that SARS-CoV −2 persistence is highly dependent on surface composition. For example, metallic surfaces (e.g., nickel silver, brass) cleared infectivity within 3 days, consistent with reports that non-porous metals accelerate viral degradation due to oxidative properties and low viral adhesion [15, 7]. Conversely, porous materials like reed and music scores retained viable virus for up to 7 days, mirroring findings on the prolonged survival of SARS-CoV-2 on such material at low level [16, 14]. In fact, SARS-CoV-2 persistence in porous materials is governed by a balance between protective entrapment (microclimate stabilization, reduced UV/desiccation exposure) and inactivating interactions (adsorption, structural disruption), leading to lower surface recoverability but potential microenvironmental survival [17, 18]. The persistence on gold-plated brass (4 days) and varnishes (e.g., shellac, linseed oil) may reflect organic coatings that shield virions from environmental stressors, as noted in studies of epoxy and polymer-coated surfaces [7, 8].

A 3-day quarantine sufficed for most non-porous materials (e.g., metals, ABS plastic), making this a feasible baseline for musicians and institutions. This is more or less in line with the recommendations adopted by the National Federation of State High School Associations (NFHS), the National Association for Music Education (NAfME) and the NAMM foundation that stated up to five days for brass and up to 3 days for plastic (https://ww1.namm.org/sites/www.namm.org/files_public/resources/COVID-19%20Instrument%20Cleaning%20Guidelines.pdf). In contrast, the recommended 4 days for instruments made of varnished wood is well-suited for shellac, linseed oil and epoxyde varnishes but overestimated for nitrocellulose- and polyurethane varnished wood. PASIC (Percussive Arts Society) guidelines recommended a 2-3 days quarantine for high-touch instruments to prevent pathogen transmission.

The extended durations required for reed (6 days) and music scores (7 days) highlight challenges for materials with complex textures or absorbent properties. These results echo studies reporting prolonged SARS-CoV-2 viability on porous surfaces [17, 18]. Notably, the inability to test ancient paper due to porosity underscores the need for alternative inactivation methods (e.g., chemical disinfection) for fragile or historically valuable items.

Experiments were conducted at 19–21°C and 50–60% relative humidity—conditions representative of indoor environments. While these parameters align with real-world scenarios, prior studies suggest that higher temperatures or humidity could accelerate viral decay [4, 6]. This raises questions about whether shorter quarantine periods might suffice in warmer climates, a variable warranting further investigation.

There are limitations in our study: (i) Laboratory conditions may not fully reflect real-world handling (e.g., frequent contact, airflow variations). Ad-hoc studies tracing viral transfer risks during instrument use post-quarantine would complete our findings; (ii) While 16 materials were tested, musical instruments often combine multiple components (e.g., wood + metal + varnish). Composite-material testing could refine guidelines; (iii) the use of a single early SARS-CoV-2 strain (UVE/SARS-CoV-2/2020/FR/702) may limit the generalizability to variants with altered environmental stability.

For the latter, despite previous studies have reported differences in the stability of the Alpha and Omicron variants [19] data on the early strains need to be considered. However, a study indicated that early variants of concern may have increased aerosol and surface stability, though there is no evidence of such an increase for the late Delta and Omicron variants [20] This is interesting because these two variants specifically caused massive waves of infection [21] Combining the experimental results with real-world data suggests that the explanation for the significant increase in infectivity and transmissibility probably lies in changes in biological properties, such as ACE-2 affinity, humoral immune escape, and mode of entry [22-25].

The results observed in this study point out (i) to investigate inactivation kinetics under varying temperature/humidity conditions, (ii) to evaluate viral transfer risks from quarantined surfaces to hands or aerosols and (iii) to explore synergies between quarantine and low-impact disinfection methods (e.g., UV-C light) for high-risk materials. Moreover, our data could be used as ground rules for other enveloped airborne viruses in period of high circulation.

The technical issue encountered with ancient scores which are fragile and frequently an historical and patrimonial resource to be protected underlines the need to anticipate such situations and to make them available through numerisation in order to maintain easy access without engaging in the risk of infectious agent transmission.

In conclusion, our work supports the adoption of stratified quarantine protocols based on material type, balancing safety with practicality [26]. For example, orchestras or schools could prioritize 3-day quarantines for metallic instruments while reserving 7-day periods for reeds and sheet music. Such guidelines could mitigate transmission risks without relying on chemical disinfectants that may damage delicate materials.

## PATENTS

### Author Contributions

Conceptualization, RNC, MJ and FB; writing and validation of original draft, BP, FT, RV, MC, JSM, FR, MJ, FB and RNC; writing—review and editing, BP, FT and RNC. All authors have read and agreed to the published version of the manuscript.

### Funding

F.T. is supported by the IRD Chair “Antiviral strategy for emergence in the South”, in partnership with Aix-Marseille University, Inserm and ANRS MIE.

